# Multi-fiber array-based photometry system for multi-regional functional mapping in the mouse brain

**DOI:** 10.64898/2025.12.23.696162

**Authors:** Manil Bradai, Mirna Merkler, Gabriela Gil, Rebecca Davie, Shuzo Sakata

## Abstract

Mesoscopic functional brain mapping is essential to better understanding of various brain functions and dysfunctions. However, accessing distributed neural circuits in mammalian brain regions remains a significant challenge. While fiber photometry is a versatile optical approach, existing methods often suffer from invasiveness and scalability. Here we present an affordable multi-fiber array (MFA)-based photometry system to monitor neural signals across multiple regions. Our system comprises a custom-designed MFA utilizing 50-µm diameter optical fibers and off-the-shelf optical components. To demonstrate the system’s versatility, we monitored GABAergic population activity using jGCaMP8s across multiple brain regions in head-fixed, awake mice. By combining with pupillometry, we identified state-dependent, region-specific GABAergic dynamics. Our MFA-based photometry system opens new avenues for investigating state-dependent neural dynamics at the mesoscopic level. To facilitate wider adoption, all codes and resources are publicly available on GitHub (https://github.com/Sakata-Lab/MFA).

## Introduction

Brain functions emerge from collective actions of distributed neuronal and non-neuronal populations (Ahrens et al., 2013, Allen et al., 2019, Atanas et al., 2023, Buzsaki, 2010, International Brain et al., 2025, Kastanenka et al., 2020, Mann et al., 2017, Oliveira and Araque, 2022, Steinmetz et al., 2019). Abnormalities in such brain-wide ensembles are associated with various brain disorders (Wang et al., 2022, Fornito et al., 2015, Noel et al., 2025, Carrera and Tononi, 2014). Therefore, it is essential to interrogate these neural and non-neural dynamics at the mesoscopic scale.

Over the past several decades, we have witnessed the explosion of various advanced technologies, allowing for monitoring of neural and non-neural population activity in an unprecedented manner (Wang et al., 2022, Machado et al., 2022). Electrophysiology remains the gold standard for monitoring individual spiking activity across the brain (Steinmetz et al., 2021, Jun et al., 2017, Buzsaki et al., 2012, Buzsaki, 2004, Angotzi et al., 2019). However, electrophysiological methods are limited in their ability to identify specific neuron types or monitor signals from non-neuronal populations, such as astrocytes.

Functional ultrasound imaging is an emerging technology that permits accessing deep brain regions (Brunner et al., 2020, Mace et al., 2011, Renaudin et al., 2022, Urban et al., 2015). However, common approaches rely on vascular dynamics rather than direct neuronal signals. The development of genetically encoded ultrasound sensors remains in the early stages compared to optical sensors (Bourdeau et al., 2018, Hurt et al., 2023, Rabut et al., 2020).

In parallel, a wide range of optical approaches have been developed, including multiphoton microscopy, light-sheet microscopy, and optoacoustic imaging (Dodt et al., 2007, Wang et al., 2003, Xu et al., 2024). Multiphoton microscopy is applicable even to freely behaving animals (Accanto et al., 2023, Klioutchnikov et al., 2023, Wu et al., 2025, Zong et al., 2022). However, while these advanced imaging techniques are robust and powerful, the scalability and affordability present significant barriers to widespread adoption.

Fiber photometry represents a versatile and accessible alternative (Byron and Sakata, 2024, Simpson et al., 2024). First introduced to systems neuroscience in 2005 (Adelsberger et al., 2005), photometry has evolved alongside the development of genetically encoded sensors (Chen et al., 2013, Marvin et al., 2013, Muir et al., 2024, Nakai et al., 2001, Patriarchi et al., 2018, Sun et al., 2018, Zhang et al., 2023) to become a standard tool. It can be combined with electrophysiology to leverage the complementary advantages of both modalities (Tsunematsu et al., 2023, Patel et al., 2020, Lewis et al., 2024, Legaria et al., 2022), and tapered fibers allow for depth-resolved monitoring in freely behaving mice (Pisano et al., 2019, Byron et al., 2025).

While conventional photometry setups utilize only one or a few optical fibers, pioneering studies have attempted to scale this up (Sych et al., 2019, Kim et al., 2016, Guo et al., 2023, Vu et al., 2024). For example, Kim et al. (2016) implanted up to seven conventional fibers, and others have adopted a multi-fiber array (MFA) with mechanical transfer (MT) ferrules to monitor over 10 brain regions simultaneously (Sych et al., 2019, Guo et al., 2023). However, these approaches suffer from limitations regarding invasiveness and flexibility. As the number of fibers increases, tissue displacement becomes a serious issue due to the large fiber diameter (≥100 µm). Furthermore, the fixed design of standard MT ferrules limits the flexibility of target coordinates. Recently, Vu et al. (2024) developed a novel MFA-based photometry system to address these limitations. While this approach was successfully deployed *within* a single targeted region (the striatum), there is a need to expand this capability to target distributed brain regions in awake mice.

Here we present an optical system designed to meet this need. Our system offers multiple features. First, the MFA design is flexible and scalable; we developed a Python script to easily customize and print a wide range of grid designs. Second, we constructed an affordable optical system using off-the-shelf components. As a proof-of-concept, we utilized this system to characterize state-dependent, region-specific GABAergic activity across multiple brain regions in head-fixed, awake mice.

## Materials and Methods

### Optical system

#### Main optical configuration

The part list is provided in **Supplementary Table 1**. In the optical system (**Fig. 1A**), 470 nm (M470L, Thorlabs) and 405 nm LEDs (M405L, Thorlabs) were used as light sources. Collimating their light (ACL2520U-A, Thorlabs), the MFA surface was illuminated through a 10x objective lens (RMS10X-PF, Olympus) to excite the fluorophores. Emission light was filtered (MF525-30, Thorlabs) and detected by a scientific complementary metal-oxide semiconductor (sCMOS) camera (ORCA-Fusion BT, Hamamatsu Photonics) (**Fig. 1B**). The sCMOS camera was connected to a computer via USB3 cable. A motorized module (PLSZ, Thorlabs) was used to adjust axial focus. The optical system was mounted on a breadboard (MB4560/M, Thorlabs). The maximum light outputs from the objective lens were 74.4 mW at 470 nm and 64.2 mW at 405 nm.

**Figure 1.**
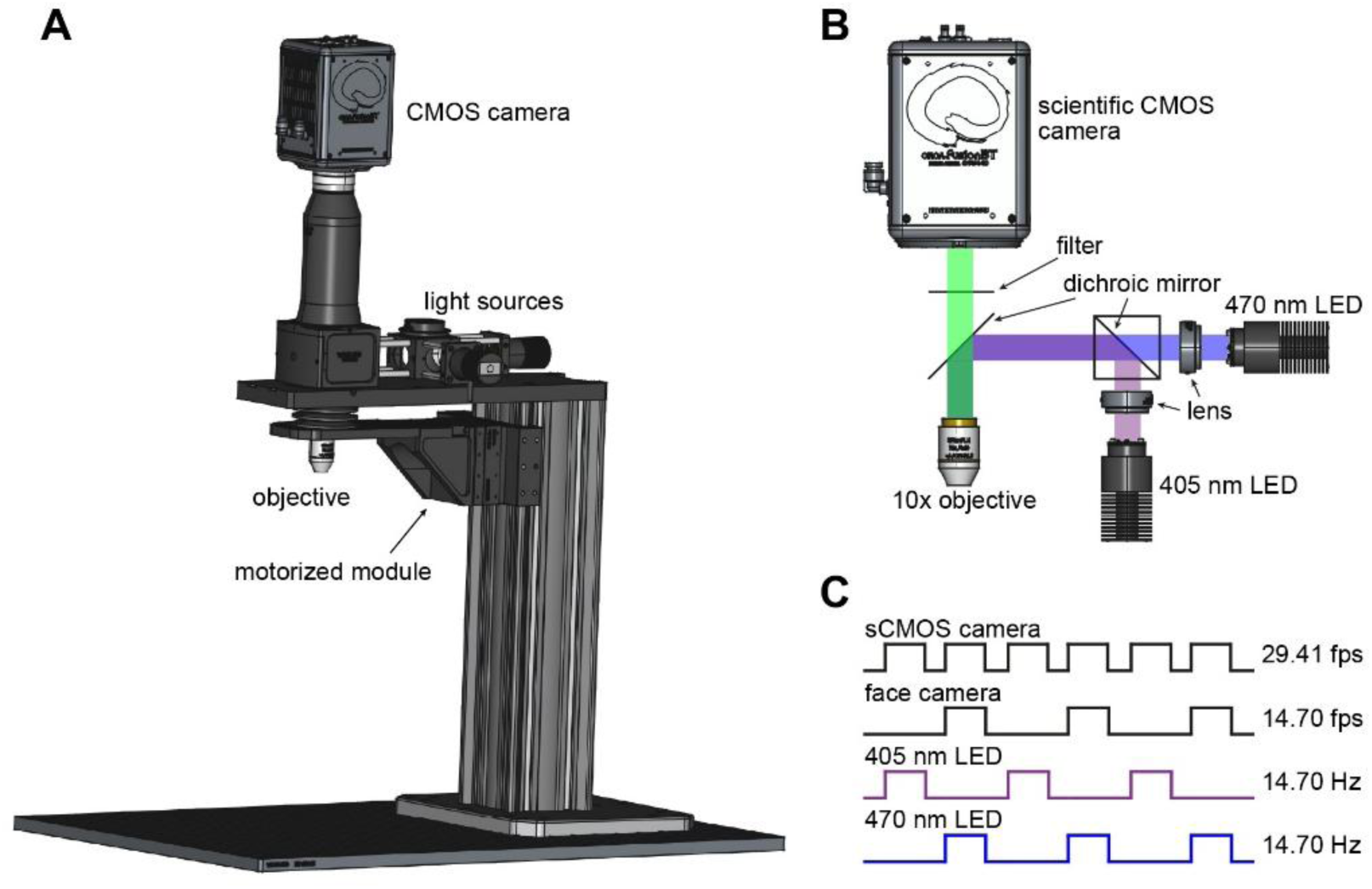
Optical system and image acquisition protocol. (**A**) A diagram of the main optical system. (**B**) A diagram of the light path. (**C**) Image acquisition protocol. Each ON pulse lasts 23 ms. Face camera images were captured when the 470 nm LED was on.

#### Peripheral devices

Each LED was driven by an LED driver with a trigger mode (LEDD1B, Thorlabs). The driver was controlled by 3.3 V analog signals generated by a data acquisition module (NI USB-6002, National Instruments). The triggering signals were gated through a custom-built OR gate (with two diodes and one register) to generate synchronization pulses for the sCMOS camera. The triggering signals for the 470 nm LED were used as synchronization signals for a face camera (aca1920-25um, Basler) connected to the imaging computer via USB3. On the breadboard, a lab jack (Labasics Lab Scissors Jack, Amazon) was mounted for coarse axial adjustment. For fine X-Y manipulation, an XY linear stage (SEM80-AS, Amazon) was used. On top of the stage, a custom-made platform (MB4, Thorlabs) with a holding tube was mounted. The face camera was placed diagonally in front of the mouse’s face. An infrared (IR) filter (FGL780, Thorlabs) was attached to the lens (M0814-MP2, Computar). An IR LED array was used as a light source.

#### Image acquisition

An image acquisition protocol is shown in **Figure 1C**. Trigger signals were controlled with a custom-written program (LabVIEW). Epifluorescent images were captured at 29.41 frames per second (fps) by triggering the sCMOS camera with LED triggering signals. HCImage Live (Hamamatsu) was used for epifluorescent image acquisition. Face camera images were captured at 14.70 fps by synchronizing them with 470 nm pulses. Pylon Live (Basler) was used for image acquisition.

### Multi-fiber array

#### Fabrication

Grids (**Fig. 2A**) were designed using CAD software (FreeCAD) and printed with high-temperature liquid (HTL) resin (IPFL). For grids with a large number of holes, a Python-based macro was used to design automatically.

**Figure 2.**
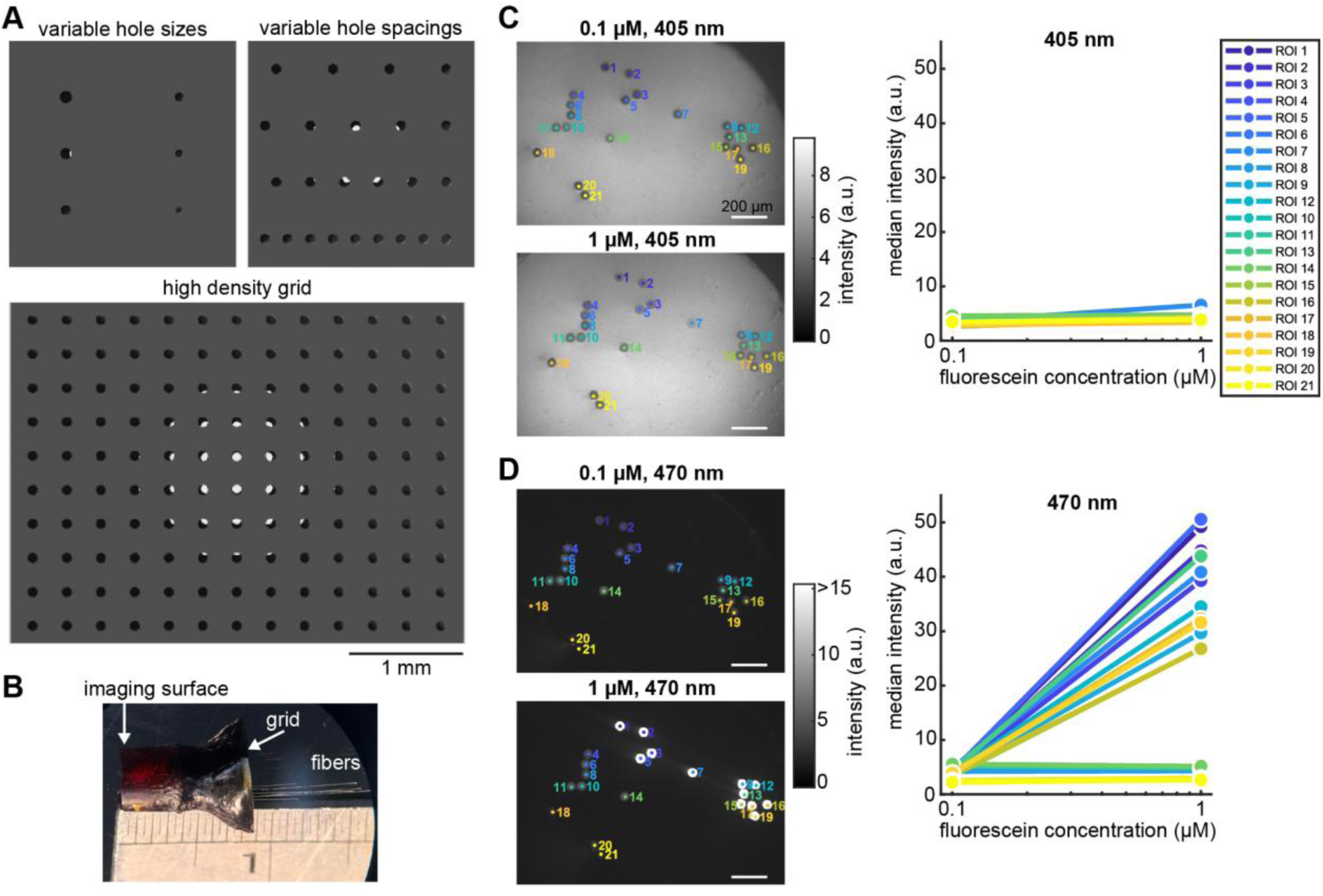
Multi-fiber array (MFA). (**A**) Custom-designed grids. (**B**) A photograph of a fabricated MFA. (**C and D**) Quality control of a MFA at 405-nm (**C**) and 470-nm LED illumination (**D**). *Left*, imaging surface with detected fibers at two fluorescein concentrations. *Right*, signal intensities across fibers as a function of fluorescein concentration. In this example, a subset of fibers was defective (no increase in signal intensity).

The detailed process of fabricating the MFA is provided in **Supplementary Information**. Briefly, each MFA (**Fig. 2B**) was fabricated using 50-µm core optical fibers (PureWave XL66, 50-µm, 0.66NA, Fiberoptics Technology Inc.), which were cut into ∼3 cm segments using a fiber scribe (S90R, Thorlabs). Each fiber was inspected under a microscope (AD106S, Andonstar) to verify a clean and flat cleave. A 3D-printed grid was secured perpendicular to the benchtop using a helping-hand and tweezers, with Blu Tack applied to one tweezer arm to improve grip and stability. Measuring and marking fiber implantation depths, fibers were inserted into the grid under a stereomicroscope (SZ51, Olympus). After insertion, the grid was rotated so that the fibers were parallel to the benchtop, and fibers were secured with black glue (RT80 Black Cyanoacrylate, RS Components).

The distal fiber ends were then gathered into a ∼1-cm segment of polyamide tubing (710-XV5, Microlumen), which was filled with superglue (ZAP THIN CA) using a microliter syringe to ensure homogeneous filling without air bubbles. Assemblies bonded with instant glue were left to dry overnight.

After consolidation of the distal bundle, the space between the polyamide tube and the grid was filled with black glue to protect the fibers. The imaging surface was then polished (30-µm, 6-µm and 3-µm polishing papers, Thorlabs).

#### Channel mapping

Fiber mapping was performed by inverting the MFA and positioning a microscope (4K WIFI Microscope, Ninyoon) beneath the imaging surface. Under a stereoscope, rows of fiber arrays were mapped first by inserting a 90°-bent piece of paper between adjacent rows to isolate the illumination. An LED pen light was directed onto the exposed row and the corresponding illuminated fibers on the imaging surface were recorded. Individual fibers within each row were subsequently identified by masking all but one fiber with a second small paper strip and illuminating the exposed fiber. The same procedure was repeated across all rows and columns until each fiber could be uniquely identified.

#### Quality control

After fabrication and mapping, each MFA was checked by immersing the fiber tips in fluorescein solution (0.1 or 1 µM in phosphate-buttered saline, PBS). As shown in **Figures 2C and D**, although no noticeable differences were detected at 405 nm illumination, signal intensities in the region-of-interests (ROIs, fibers) were increased as the fluorescein concentration increased if the fibers were intact.

### Animals

All animal experiments were performed in accordance with the United Kingdom Animals (Scientific Procedures) Act of 1986 Home Office regulations and approved by the University of Strathclyde Animal Welfare and Ethical Review Body and the Home Office (PPL0688944). PV-IRES-Cre (PV-Cre; JAX008069) or GAD2-IRES-Cre mice (GAD2-Cre; JAX010802) were bred with TIGRE2-jGCaMP8s-IRES-tTA2-WPRE mice (jGCaMP8s; JAX037952) on the C57BL/6J background. All genotyping was performed by Transnetyx using real-time PCR. Two male PV-Cre;jGCaMP8s (34 and 40 weeks old) and two male GAD2-Cre;jGCaMP8s (30 and 32 weeks old) mice were used for MFA-based photometry experiments. They had *ad libitum* access to food and water. Until surgery, the animals were housed with their littermates on a 12/12 h light/dark cycle. All imaging sessions were performed at zeitgeber time 5-7.

### Surgery

Key surgical variables are listed in **Supplementary Table 2**. Mice were anesthetized with isoflurane (1%–1.5%) and placed in a stereotaxic apparatus (DKI 961, Kopf). To provide analgesia, Ropivacaine (Naropin, 8 mg/kg) was administered subcutaneously at the site of the incision, while Carprofen (Rimadyl, 20 mg/kg), Buprenorphine (Vetergesic, 0.1 mg/kg) and Dexamethasone (Rapidexon, 0.2 mg/kg) were administered subcutaneously at the back. The skin was removed from the planned head cap area, and the periosteum was cut, scraped, and dissolved with 3% hydrogen peroxide. Three anchor screws (418–7123, RS Components) were implanted into the skull. Two screws were placed in the front at coordinates AP -0.5 mm, ML +1.5 mm and AP -1.5 mm, ML -2.5 mm in one mouse, or at AP -1.5 mm, ML -1.5 mm and AP -3.0 mm, ML -4.0 mm in three mice. The third screw was placed on the cerebellum (AP - 6.0 mm, ML +2.0 mm). A custom-made semicircular headpost was attached to the skull anteriorly. A medium-size craniotomy was performed above the right hemisphere (AP –3.5 to -5.5 mm, ML 0 to +2 mm; or AP –0.5 to -2.5 mm, ML +0.5 to +2.5 mm; or AP -0.5 to -4.5 mm, ML 0 to +3.5 mm). MFAs were implanted. In GAD2-Cre;jGCaMP8s mice, a durotomy was performed before implanting the MFA. The MFA was implanted at maximal depths of 1.35 mm (n=1), 4.1 mm (n=1) or 4.5 mm (n=2), measured from the brain surface. After implantation, the exposed brain surface was covered with a biocompatible sealant (Kwik-Sil, World Precision Instruments). Layers of dental cement and cyanoacrylate were used to cover the skull screws and skull surface, as well as to secure the headpost and MFA in place. A flat-lid Eppendorf tube was cut to the desired height and secured over the MFA for protection. The final layer of dental cement was mixed with black ink to create a black headcap. After surgery, mice were housed in high-topped cages with *ad libitum* access to food and water, and allowed to recover for at least 5 days.

### Imaging procedures

For habituation to a head-fixed condition, the animals were secured in the head fixation apparatus through the headpost and placed into an acrylic tube layered with absorbent paper. Habituation lasted 3-5 days, during which the head fixation duration was gradually extended from 15 to 60 minutes.

For imaging sessions, after placing an animal in the head-fixation apparatus described above, the MFA imaging surface was focused under the microscopic imaging system in a dim light condition. An imaging session lasted 10-30 minutes while monitoring the mouse’s face. The exposure time was set 22 ms. LED power was set to 26.8∼64.9 mW at 470 nm and 2.8∼14.5 mW at 405 nm.

### Perfusion

After completing imaging sessions, the mice were deeply anesthetized with pentobarbital (200 mg/ml) and transcardially perfused with PBS and 4% paraformaldehyde. After cutting the neck, the lower jawbone and surrounding tissues were removed to maximize access to a contrast reagent for micro-computed tomography (µCT) scanning (see below) while maintaining the brain and MFA intact. The sample was incubated in the same fixative for at least 12 hours at 4°C. The next day, after washing the sample with PBS for 5 min three times, the sample was incubated in Lugol’s solution (2.5% potassium iodine, 1.25% iodine in distilled water) at room temperature for 6-10 days.

### µCT scan

After washing the sample with PBS for 5 min three times, µCT images were taken (Quantum GX2, PerkinElmer). The sample was scanned at 90 kV and 88 µA. The field of view (FOV) was set to 36 mm for acquisition and 10 mm for reconstruction (10 µm voxel size). The scan mode was set to “high resolution” mode with 4 min scanning.

### Data analysis

The experimental overview and data analysis pipelines are summarized in **Figure 3**. In each animal, three types of data were acquired (**Fig. 3B**): MFA-based photometry data, face camera images, and µCT images.

**Figure 3.**
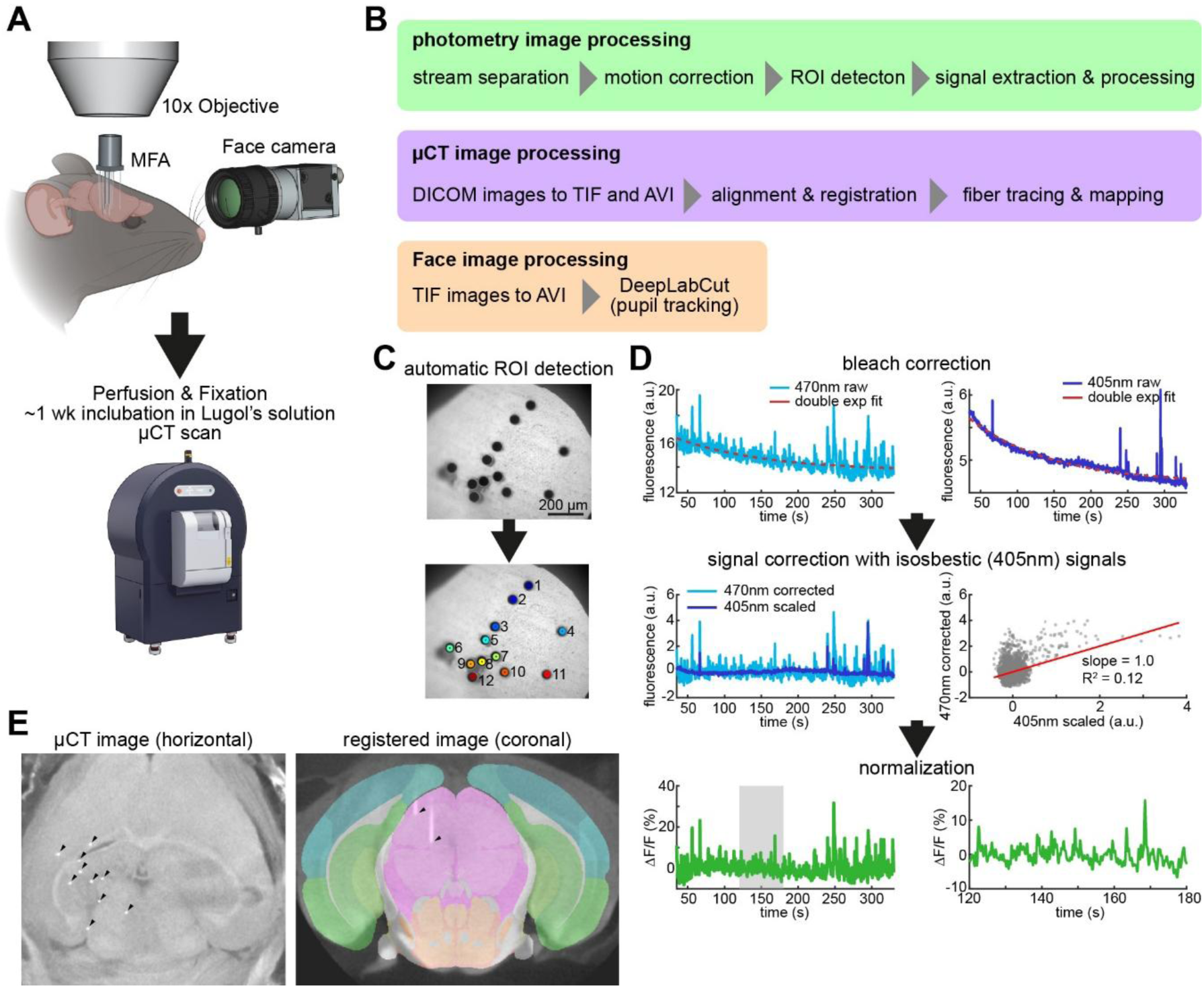
Data analysis workflow for MFA-based photometry experiments. (**A**) Diagram outlining an experiment. After *in vivo* photometry experiments, the mouse head was scanned with a µCT scanner. Images (mouse and µCT scanner) were created by bioRender and Gemini 2.5 Pro. (**B**) Data processing workflows for three types of images: photometry images (top), µCT scan images (middle) and face images (bottom). (**C**) An example of automatic region-of-interest (ROI) detection. Top, raw 405 nm image. Bottom, image with detected ROIs (fibers). (**D**) Photometry signal processing. Extracted fluorescent signals from each ROI were processed based on this workflow. Top, photobleach correction with a double-exponential curve. Middle, signal correction with scaled isosbestic (405 nm) signals and correlation between isosbestic and 470 nm signals. Bottom, normalization (z-scoring). The right panel shows stretched signals in the shaded time segment of the left panel. (**E**) µCT scan to localize fiber positions. Left, horizontal image with multiple fibers (arrow heads). Right, coronal image with two fibers (arrow heads) registered to the Allen CCFv3.

#### Photometry image processing

Image processing was performed offline using a custom-written MATLAB script (R2024a, MathWorks).

After loading a .cxd file, time stamps were extracted to assess frame drops. Based on the extracted time stamps, the image streams of 405 nm and 470 nm illumination were separated and each image stream was saved separately. When a frame drop was detected, previously available frames were used. After stream separation, artificial motion was corrected using NoRMCorre (Pnevmatikakis and Giovannucci, 2017). Since 405 nm images were often underperformed due to high autofluorescence at the imaging surface, pre-processing (a subtraction of a Gaussian blur from the original image) was performed before motion correction.

To detect fibers (ROIs), a 405 nm frame was processed to detect circles (MATLAB’s imfindcircles.m) and the ROI number was assigned automatically (**Fig. 3C**). Each ROI was defined as a circle with a 2-pixel diameter. The median intensity of each ROI in each frame was computed to establish raw fluorescent signals. Based on these fluorescent signals, a conventional photometry signal processing workflow was applied (**Fig. 3D**) (Patel et al., 2020, Simpson et al., 2024): (i) based on a double-exponential curve, photobleaching was estimated and corrected. (ii) motion artifacts were corrected based on scaled 405 nm (isosbestic) signals using linear regression. (iii) ΔF/F was defined as normalized (Z-score) motion corrected signals.

#### Face image processing

Since the face camera captured multiple .tiff files, first, a movie file (.avi) was constructed. Using DeepLabCut (version 3) (Mathis et al., 2018), the four edges (top, right, bottom, and left) of the pupil were tracked. The smoothed profile was used for further analysis. The likelihood threshold was set to 0.6. The pupil diameter was estimated as the mean of vertical and horizontal pupil diameters. Low-pass (0.2 Hz) filtered signals were used for further analysis.

#### µCT image analysis

Image processing (**Fig. 3B**) was performed using a custom-written Python package (mfaCTpy, https://github.com/Sakata-Lab/mfaCTpy).

After constructing a .tif file from .dcm files, the volumetric image was aligned with the midline by manually labeling the midline on multiple horizontal sections. The aligned image was registered to the Allen Common Coordinate Framework (Wang et al., 2020) by manually labeling multiple landmarks (Sergejeva et al., 2015). Each fiber was traced by manually detecting the fiber tip in the brain and the top near the grid. This positional information was aligned with the fiber mapping information obtained during the MFA fabrication (see above). **Figure 3E** shows an example of a horizontal µCT image and a registered coronal image.

#### Photometry-pupil correlation analysis

Arousal states were classified based on normalized (Z-score) pupil diameter signals. States were separated into three states (high, medium, and low arousal) using quantiles (33% and 67%).

For MFA-based photometry signals, fibers providing calcium transients were selected and only a time window without excessive artifacts was used. Processed photometry signals were bandpass (0.5 – 1 Hz) filtered, and power was estimated by computing the moving variance with a -5-sec sliding window. Then the power was normalized (Z-score). To assess the correlation between pupil and photometry signals, Pearson’s correlation coefficient was computed.

### Code repository

Codes and resources are available on GitHub (https://github.com/Sakata-Lab/MFA).

## Results

Although GABAergic neurons are a major cell class in the brain (Yao et al., 2023), functional mapping of GABAergic neural populations across multiple brain regions remains challenging. Additionally, although parvalbumin-positive (PV+) neurons, a subset of GABAergic neurons, have been extensively investigated in the isocortex and hippocampus, less is known about PV+ neurons in the subcortical regions. We hypothesize that our MFA-based photometry system is suitable for this technical challenge. As a proof-of-concept, we monitored GABAergic (GAD2+) or PV+ neurons across multiple subcortical regions in head-fixed awake mice (**Fig. 4**). We used two transgenic lines, GAD2-Cre;jGCaMP8s (2 males) and PV-Cre;jGCaMP8s (2 males) while monitoring the pupil as a biomarker of arousal. A subset of fibers provided calcium transients (**Supplementary Table 3**).

**Figure 4.**
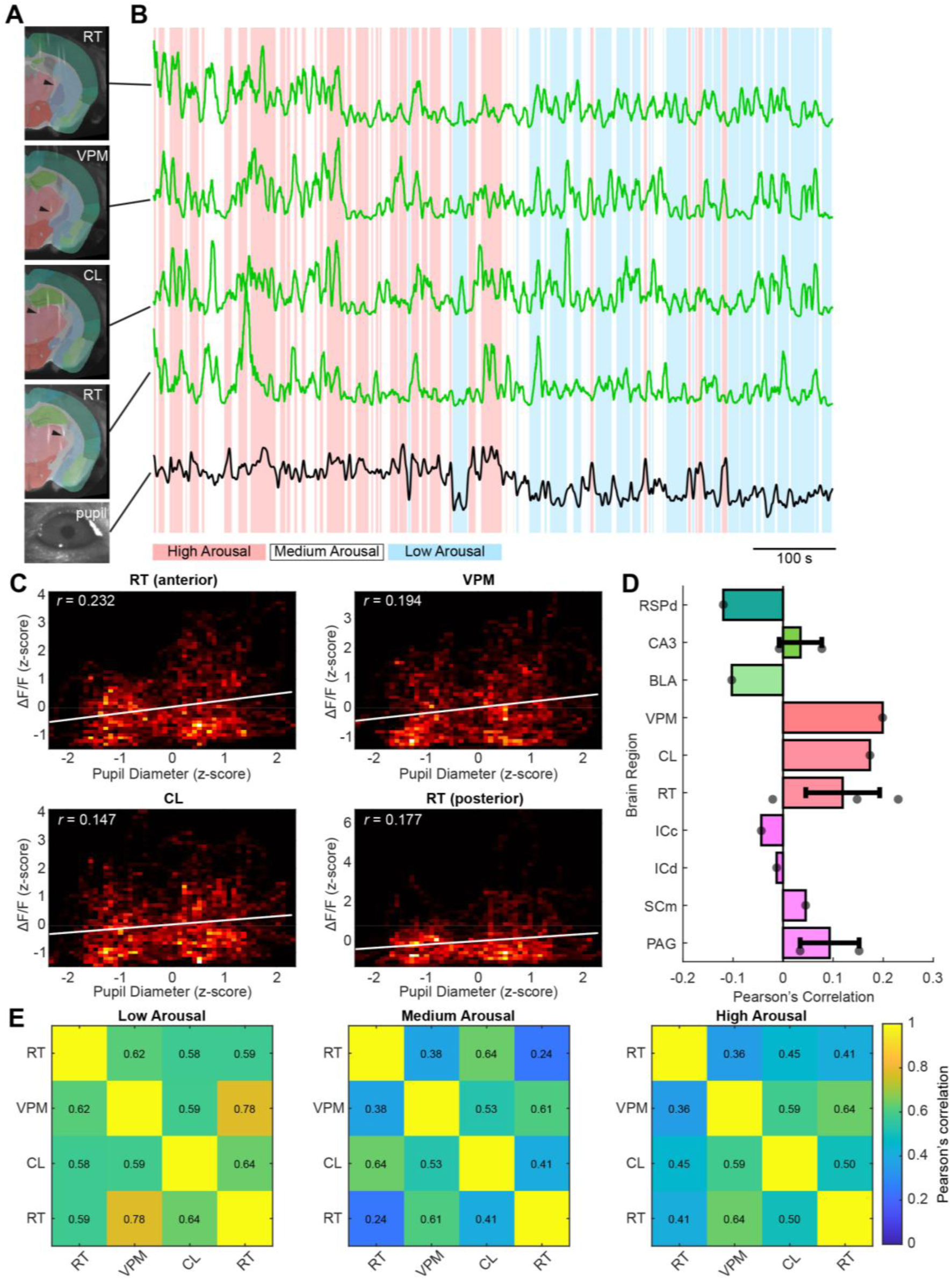
MFA-based photometry to investigate state-dependent, region-specific population activities. (**A**) µCT images with fiber tips (arrow heads) registered to the Allen CCF. Bottom, a mouse eye image. RT, reticular nucleus of the thalamus. VPM, ventral posteromedial nucleus of the thalamus. CL, central lateral nucleus of the thalamus. (**B**) Sample photometry traces across four subcortical regions with pupil dynamics. Background colors indicate different arousal states based on pupil size. (**C**) Density heatmap of pupil-photometry signal correlation across brain regions. Solid line, linear regression line. *r*, Pearson’s correlation coefficient. (**D**) Summary of Pearson’s correlation coefficient across the four experiments. RSPd, retrosplenial area, dorsal part. BLA, basolateral amygdalar nucleus. ICc, inferior colliculus, central nucleus. ICd, inferior colliculus, dorsal nucleus. SCm, superior colliculus, motor-related. PAG, periaqueductal gray. Error bars indicate the standard deviation. (**E**) An example of functional connectivity maps (correlation matrix) across three arousal states.

With this dataset, we began by investigating signals in a PV-Cre;jGCaMP8s mouse (**Figs. 4A and B**). Using a µCT scanner, we confirmed that fibers were located in the anterior reticular thalamus (RT), ventral posteromedial nucleus of the thalamus (VPM), central lateral nucleus of the thalamus (CL) and the posterior RT (**Fig. 4A**). Calcium transients appeared densely and rhythmically probably because we monitored population activity in PV+ neurons. Since we noticed that 0.5-1 Hz band signals reflect the rhythmic nature of their population activities (**Supplementary Figure 1**), we estimated the power of bandpass-filtered signals across channels (**Fig. 4B**). When we compared photometry signals with pupil diameter, we noticed correlations between them (**Fig. 4C**). Across four experiments, there was a tendency that the correlations between photometry signals and pupil size are variable depending on brain regions (**Fig. 4D**).

Finally, since we monitored calcium signals across multiple brain regions simultaneously, we examined if functional connectivity between brain regions changed with arousal. Since pupil size reflects arousal, we classified pupil size into three states: high, medium and low arousal states (**Fig. 4B**) (**Supplementary Figure 2**). Based on this classification, we observed that functional connectivity between brain regions changes across arousal states (**Fig. 4E**). Overall, the MFA-based photometry system allows us to monitor cell-type-specific population activity across multiple subcortical regions and arousal states.

## Discussion

Brain-wide cell-type-specific functional mapping is a major technological challenge in modern systems neuroscience. In this study, we presented a MFA-based photometry system that allows us to monitor neural population activity across multiple brain regions and brain states in a head-fixed condition. Our photometry system expands the existing toolbox to open new avenues for the investigation of brain-wide neural ensembles across cell-types and brain states.

### Comparisons with existing photometry systems

Fiber photometry is a widely adopted optical approach to monitoring various signals in the brain. Although conventional fiber photometry uses a thick (200∼400 µm diameter) optic fiber, our system can accommodate multiple thin (50 µm diameter) fibers, which targets distributed neural circuits even in deep brain regions.

A few studies have pioneered photometry with multiple fibers (Vu et al., 2024, Sych et al., 2019, Kim et al., 2016). Kim et al. (2016) implanted up to 7 conventional optic fibers simultaneously whereas Sych et al. (2019) utilized MT ferrule technology with up to 48 fibers. Since the diameter of their optic fibers was 100-400 µm, the tissue displacement is 4-64 times larger than a 50-µm diameter fiber. Another major difference is the flexibility of MFA design (**Fig. 2A**). By designing a grid according to experimental requirements, our approach allows us to target distributed brain regions with minimal invasiveness.

Our system was inspired by Vu et al. (2024). Differences from their original system can be summarized as follows: first, our light sources are low-profile while we achieved sufficient light outputs for photometry by optimizing the light path configuration with collimators. This makes our system more affordable for the wider community. Second, we demonstrated neural signals across multiple brain regions simultaneously while the original system targeted only the striatum. Third, we optimized data analysis workflows. For example, knowing that a 405 nm image provides high contrast between fibers and the remaining image surface, we implemented simple automatic ROI detection to extract fluorescent signals across fibers. This refinement improves scalability and reproducibility. Additionally, we developed a Python-based package to trace fibers. Since the existing approach still requires manual curation and the outcome depends on CT image quality, we decided to use simple manual tracing. We implemented a graphical user interface to facilitate this manual tracing.

### State-dependent, region-specific GABAergic activity

As a proof-of-concept, we applied our MFA-based photometry system to monitor GABAergic population activity across multiple brain regions and arousal states. Our preliminary results imply state-dependent, region-specific GABAergic neural activity. While GABAergic neurons have long been implicated in the regulation of global brain states, including sleep-wake cycles (Sulaman et al., 2023), their coordinated actions remain underexplored. Our photometry system will help address this fundamental challenge.

### Potential applications

Given the versatility of fiber photometry in general (Simpson et al., 2024, Byron and Sakata, 2024), the following applications may be considered in the future: first, it would be interesting to apply our system to monitor various signals, including astrocytic calcium signals (Tsunematsu et al., 2021, Noh et al., 2023), neurotransmitters/neuropeptides (Muir et al., 2024, Kagiampaki et al., 2023, Duffet et al., 2022), disease pathology (Byron et al., 2025) and intracellular molecular signals (Chen et al., 2013). For example, while region-specific and state-dependent calcium dynamics were examined in astrocytes (Tsunematsu et al., 2021), simultaneous monitoring of astrocytic calcium signals across the brain has not been realized. Additionally, while depth-resolved amyloid plaque signals were recently monitored in an Alzheimer’s disease mouse model by utilizing tapered fibers (Byron et al., 2025), our system may offer an alternative solution to monitoring disease pathological signals across multiple brain regions.

Second, the combination with other modalities, such as electrophysiology, will complement their advantages with each other (Patel et al., 2020, Lewis et al., 2024). For example, it would be interesting to determine how sub-second neural events, such as sharp-wave ripples and pontine waves, are associated with cell-type-specific population activity across the brain. This has not been realized by existing technology.

Third, all-optical approaches are an attractive option. Indeed, Vu et al. (2024) demonstrated the feasibility of combining optogenetic stimulation with a digital micromirror device (DMD). A potential alternative to our approach, because fibers are sparsely arranged, is to use expression-targeting approaches (i.e., to express opsins in a single brain region) by adding an additional LED.

Fourth, experiments in a freely behaving condition are another key area for future development. While a combination with a miniaturized endoscope is a solution (Vu et al., 2024), a coherent fiber bundle may also be considered to utilize the same optical setup (Accanto et al., 2023).

Finally, implementing real-time image processing would be a promising area since motion correction can be performed in real-time (Pnevmatikakis and Giovannucci, 2017). This will allow closed-loop experiments. Overall, although the present study focused on the initial development of the MFA-based photometry system, there is a wide range of potential applications and developments in the future.

### Limitations of the study

Although we successfully developed a MFA-based photometry system, there are multiple limitations to overcome. First, our MFA design and fabrication can be optimized. For example, we still need to explore the upper limit of fibers. We also noticed that a large fraction of fibers did not provide calcium signals partly due to technical errors (e.g., broken fibers, inconsistent fiber quality, mistargeted fibers, etc). The materials and design (e.g., hole diameter) of the grid also need to be optimized. Although we used yellow-translucent HTL resin in this project, carbon black resin may also be explored to reduce light penetration.

Second, our current system allows only head-fixed experiments that perturb naturalistic behavior. However, as discussed above, it would be interesting to apply a miniaturized endoscope or coherent fiber bundle in the future.

Third, the temporal resolution of our system is less than 30 fps. However, a recent study conducted fiber photometry with genetically encoded voltage indicators at 100 Hz (Haziza et al., 2025). As an advanced CMOS camera is developed, this limitation can be overcome.

Finally, while our fiber tracing relies on µCT, the poor compatibility of Lugol’s treated samples with conventional histological analyses may limit their application. Although Lugol’s treatment enhances µCT image contrast, it increases tissue rigidity and reduces accessibility to antigens for immunostaining. This will be particularly problematic if viral approaches are taken since expression patterns must be examined in every sample. In this project, we took a transgenic approach to mitigate this. While a combination of tissue clearing and advanced microscopy may be considered, MFA removal must be done without tissue damage.

## Conclusion

Mesoscopic functional brain mapping is the key to understanding brain function and dysfunction. Our MFA-based photometry system offers an innovative solution to targeting distributed neural circuits at depth. Our system contributes to the ongoing effort towards the brain-wide mapping and manipulation of neuronal and non-neuronal activity *in vivo*.

## Author contributions

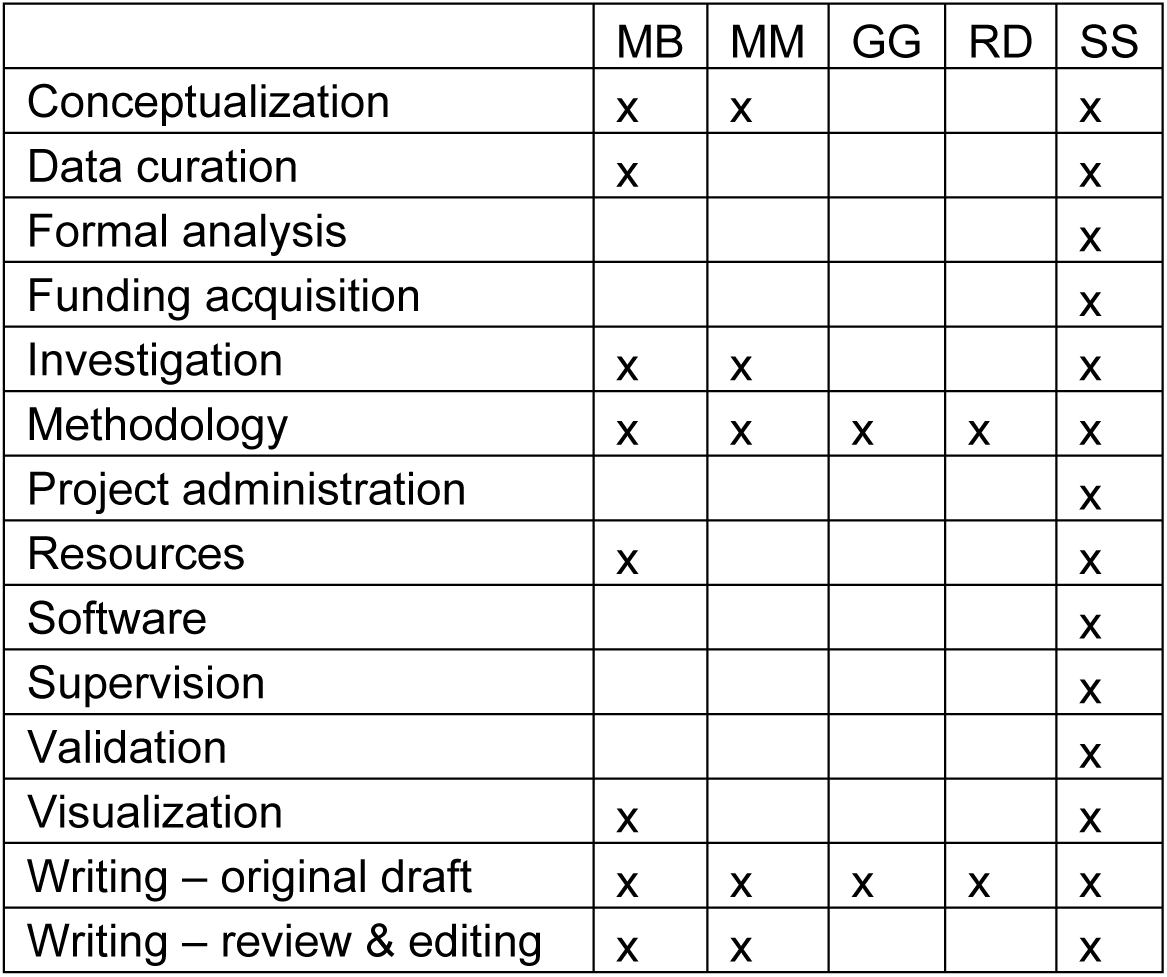

## Acknowledgements

This work was supported by Medical Research Council (MR/Y004051/1 to S.S.), and the European Union’s Horizon 2020 (H2020-ICT, DEEPER, 101016787 to S.S.).

