## Supplementary Information for "Multi-fiber array-based photometry system for multi-regional functional mapping in the mouse brain"

**Supplementary Table 1. Optical system parts list**

| product name | product number | qty | supplier |
|---|---|---|---|
| CMOS Camera | | | |
| ORCA-Fusion BT | C15440-20UP | 1 | Hamamatsu Photonics |
| Main Body | | | |
| Camera Port | WFA4100 | 1 | Thorlabs |
| Illumination Cube Housing | WFA2002 | 1 | |
| Microscopy Cube Assemly | MDFM-MF2 | 1 | |
| Dichroic Filter | MD498 | 1 | |
| Emission Filter | MF525-39 | 1 | |
| Cerna Body with Epi Arm | CEA1400 | 1 | |
| Aluminium Breadboard | MB4560/M | 1 | |
| Objective, Arm, Motorized Module | | | |
| 10x Fluorite Objective | RMS10X-PF | 1 | Olympus |
| Objective Arm | CSN100 | 1 | Thorlabs |
| Adaptor | M32RMSS | 1 | |
| Angle Bracket | PLSZ1 | 1 | |
| Motorized Module | PLSZ | 1 | |
| Stepper Controller | MCM301 | 1 | |
| USB Joystick | MCMK3 | 1 | |
| Light Sources | | | |
| 405 nm LED | M405L | 1 | Thorlabs |
| 470 nm LED | M470L | 1 | |
| LED Holder | CP12 | 2 | |
| LED Driver | LEDD1B | 2 | |
| Aspheric Condenser Lens | ACL2520U-A | 2 | |
| Lens Tube | SM1L05 | 2 | |
| Cage Plate | CP08/M | 2 | |
| Filter Holder | C4W | 1 | |
| Kinematic Cage Cube Platform | B4C/M | 1 | |
| Dichroic Mirror | DMLP425R | 1 | |
| Peripheral (face camera) | | | |
| Camera | aca1920-25um | 1 | Basler |
| Zoom Lens | M0814-MP2 | 1 | Computar |
| IR Filter | FGL780 | 1 | Thorlabs |
| IR LED Array | USB100W05MT-DL36 | 1 | ELP |
| Peripheral (DAQ) | | | |
| DAQ Module | NI-USB-6002 | 1 | National Instruments |
| OR Gate | custom-made | 1 | RS components |
| Computer | Precision 5860 Tower | 1 | Dell |

#### Supplementary Table 2. Surgical parameters

| | | Mouse #1 | Mouse #2 | Mouse #3 | Mouse #4 |
|---|---|---|---|---|---|
| Genotype | | PV-Cre;jGCaMP8s | PV-Cre;jGCaMP8s | GAD2-Cre;jGCaMP8s | GAD2-Cre;jGCaMP8s |
| Sex | | male | male | male | male |
| Age | | 8 mo | 9 mo | 7 mo | 7 mo |
| Fiber # | | 11 | 16 | 15 | 18 |
| Craniotomy coordinates | AP | -3.5:-5.5 mm | -0.5:-2.5 mm | -0.5:-4.5 mm | -0.5:-4.5 mm |
| | ML | 0:+2 mm | +0.5:+2.5 mm | 0:+3.5 mm | 0:+3.5 mm |
| Durotomy | | No | No | Yes | Yes |
| Penetration depth | | 1.35 mm | 4.1 mm | 4.5 mm | 4.5 mm |

#### Supplementary Table 3. Parameters for data analysis

| | Mouse #1 | Mouse #2 | Mouse #3 | Mouse #4 |
|---|---|---|---|---|
| Genotype | PV-Cre;jGCaMP8s | PV-Cre;jGCaMP8s | GAD2-Cre;jGCaMP8s | GAD2-Cre;jGCaMP8s |
| Functional fiber # | 2/11 | 4/16 | 4/15 | 4/18 |
| Duration (s) | 380 | 820 | 903 | 360 |

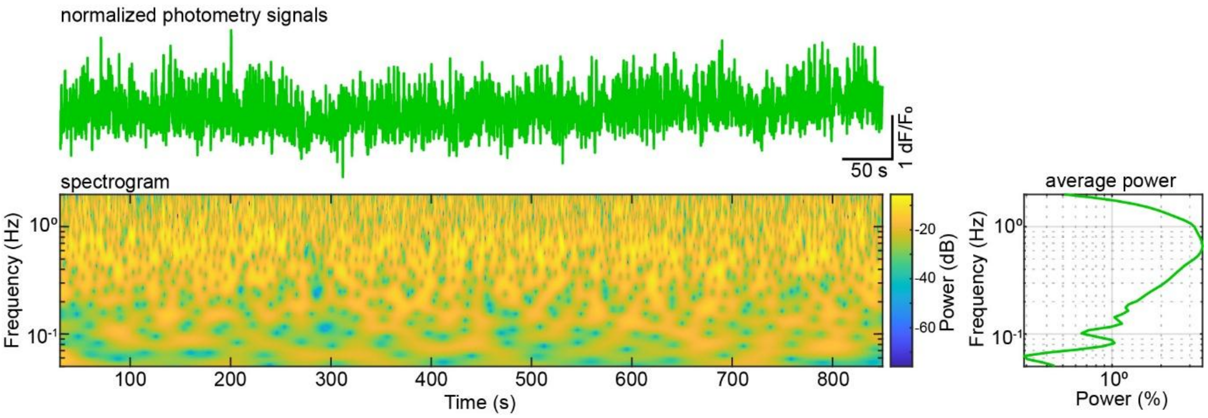

**Supplementary Figure 1. Time-frequency profile of photometry signals.** *Top*, normalized photometry signals. *Bottom left*, spectrogram of photometry signals. *Bottom right*, average power profile, normalized to percentage.

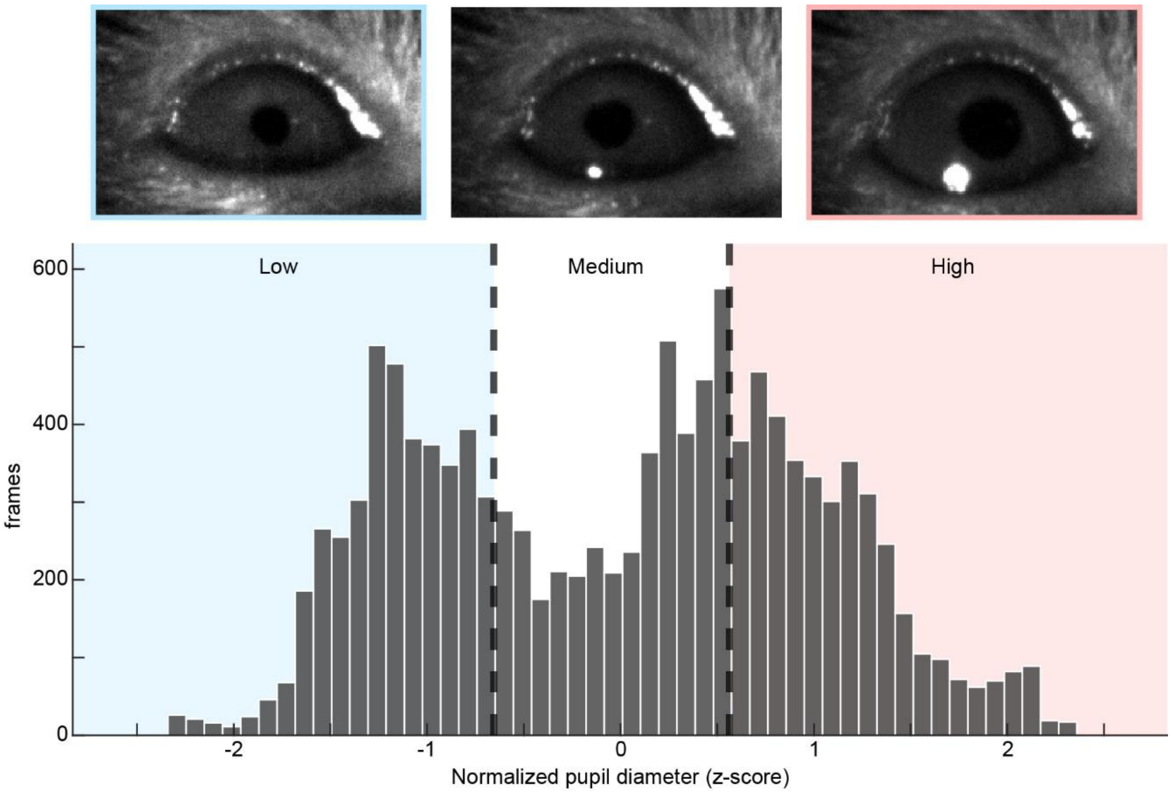

**Supplementary Figure 2. Arousal state classification based on pupil diameter**. *Top*, representative pupil images. *Bottom*, the distribution of normalized pupil diameters. 33 and 67 percentiles were the thresholds to classify arousal states.

### Multi-Fiber Array Construction Manual

Gabriela Gil and Rebecca Davie

### 1. Materials and Equipment

#### 1.1 Consumables

| Item | Manufacturer | Product Information |
|---|---|---|
| Optic fibers | Fiberoptics Technology Inc | Purewave XL66 1mmx30 Raw Fibre Bundle, 50µm, NA66 |
| MFA grid | IPFL | Custom design |
| UV glue | Norland Products, Inc | Norland Optical Adhesive, NOA61, 1 OZ, 28.5g, LOT 612 |
| Black Glue | RS Components | RS Pro, RT80 Black Cyanoacrylate, 20g, RS 908-2802 |
| Pink Zap Glue | ZAP-A-GAP | ZAP CA Thin CA, 1 OZ (28.3g), PT-08 |
| Solder Wire | MBO (UK) Ltd. | Solder Wire, 500g, 547-287 |
| Acrylic Resin Liquid | Kemdent | Simplex Rapid Acrylic Liquid, 150ml, LOT 47557 |
| Polishing Paper (green) | Thorlabs | 6x6 Inch, Diamond Lapping (Polishing) Sheets, 30µm Grit, LF30D |
| Polishing Paper (yellow) | Thorlabs | 6x6 Inch, Diamond Lapping (Polishing) Sheets, 6µm Grit, LF6D |
| Polishing Paper (pink) | Thorlabs | 6x6 Inch, Diamond Lapping (Polishing) Sheets, 3µm Grit, LF3D |
| Tubing | MicroLumen | Polyimide medical tubing, ID: 0.0710, OD:0.0988, Wall: 0.01390, 710-XV.5 |
| Tubing | MicroLumen | Polyimide medical tubing, ID: 0.0477, OD:0.0578, Wall: 0.00505, 475-V.25 |
| Reusable adhesive | Bostik | Blu Tack |
| Fluorescein HITC | Thermo Scientific Chemicals | Fluorescein Pure, 119240250 |

#### 1.2 Tools and Instruments

| Item | Manufacturer | Product Information |
|---|---|---|
| Optic fiber scribe | Thorlabs | FiberScribe, S90R |
| Stereoscope | Olympus | SZ51 |
| Fiber Polishing Pad | Thorlabs | Silicone Polishing Pad, 50 duometer, 8.75x13 Inch (222.3mmx320.2mm) and 0.12Inch Thick (3.1m), NRS913A |
| Glass Polishing Plate | Thorlabs | Glass Polishing Plate, 9.5x13.5Inch and 0.5Inch Height, CTG913 |
| LED Pen | Cakkone | CAT 2025 2 8 |
| Digital Microscope | Ninyoon | 4K WIFI Microscope, 748521049947 |
| Digital Microscope | Andonstar Technology Co.,Ltd. | AD106S |
| Tweezers | World Precision Instruments | Dumont Tweezers 5/45 |
| Tweezers | TOWOT | Precision Tweezers, 5pcs Machine Tweezers Made of Stainless Steel Heat Resistant, SA13 |
| Tweezers | Fine Science Tools | Graefe Iris Forceps, 11064-07 |
| Scissors | World Precision Instruments | Operating Scissors 12/21, 501754-G |
| Optical system | See main text | |

### 2. Designing a Multi-Fiber Array (MFA)

This section outlines how to plan and design a customized MFA by identifying target brain regions, determining their anatomical coordinates using brain atlases, and arranging fibers on a spatial grid.

- Identify the target brain regions for optical recordings or stimulation.
- Use the Paxinos and Franklin Mouse Brain Atlas or the Allen Mouse Brain Atlas to determine the anatomical location of each region of interest.
- Define the MFA grid layout (e.g., 4 × 3 mm).
  - Grid dimensions are fully customizable and should be adapted to the spatial distribution of the target regions.
  - The spacing used here was 0.3 mm, resulting in linear offsets of 0.3 mm, 0.6 mm, 0.9 mm, etc.
  - *Note that spacing is optional and can be modified depending on experimental requirements.*
- Map each fiber position on the grid to its corresponding anatomical target based on atlas reference.
- Adjust fiber lengths and array geometry to ensure accurate implantation and compatibility with the planned recording or stimulation approach.

### 3. Preparation of Optical Fibers

This section covers how to set up a clean, organized workspace and prepare individual optical fibers for later assembly. It includes cutting, inspecting, and selecting viable fiber segments that will form the implantation array.

#### 3.1 Workspace Preparation

- Lay out the Thorlabs silicone polishing pad on the Thorlabs glass polishing plate
- Spray the surface with antiseptic spray and wipe to clear any residue.
- Turn on the Andonstar digital microscope, prepare the Thorlabs fiber scribe, ruler and the Purewave fiber bundle.

#### 3.2 Cutting and Organising Viable Fibers

- Pull out a fiber (be cautious of the glass) from the Purewave fiber bundle
- Place the fiber on the silicone polishing pad horizontally,
- With a ruler measure out approximate length of 3cm (anything longer can make the later stages harder) (**Fig. 1A**).
- Cut 3cm fiber segments by aligning the Thorlabs fiber scribe perpendicular to the fiber and gently slice in a one smooth motion
  - *Note that any excess pressure or pushing the blade directly into the fiber will result in damage.*
- Inspect each of the cut fibers by holding them in tweezers no. 5/45 with a little bit of blu tack for better grip under the Andonstar digital microscope.
- Place suitable ends of the fibers facing one direction (these will be the ones used for implantation) (**Fig. 1B**)

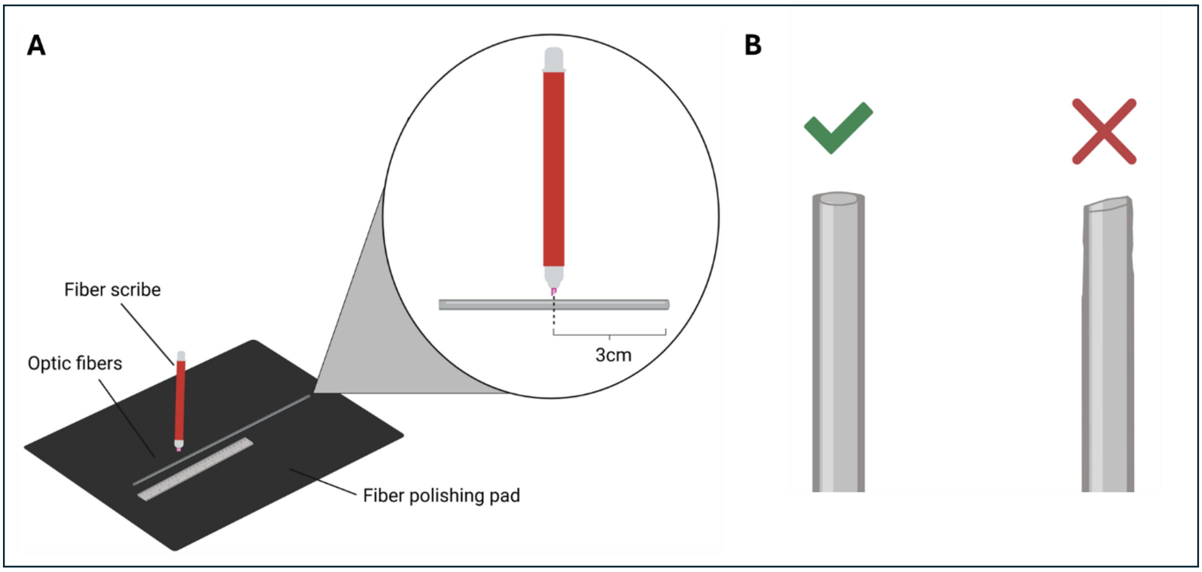

**Figure 1 | Preparation and Cutting of Optic Fibers.** Schematic illustrating the procedure for cutting optic fibers. (A) A Thorlabs fiber scribe is positioned perpendicular to an optic fiber taken from the Purewave fiber bundle. The scribe is used to cut the fiber, producing a ~3cm segment. (B) Comparison of correct (left) and incorrect (right) fiber preparations following cutting. The left image shows a cleanly cut fiber with a smooth, flat imaging surface suitable for implantation. The right image shows a poorly cut fiber with a damaged, angled imaging surface that is unsuitable for implantation. Figures not to scale. Created in bioRender.

### 4. Preparing and Mounting the Grid

Explains how to position and stabilise the MFA grid so that its holes are clearly visible and accessible. Proper mounting is essential for successful fiber insertion in the next steps.

- Take the MFA grid (depending on the flexibility of the grid find the best way to mount it) and gently close inside of the tweezer no. SA13 securing vertically in the helping hand (**Fig. 2**)
    - *Note: Apply a thin layer of blue tack to inside to better grip*
- Adjust the tilt of the grid under the Olympus Stereoscope, the holes of the grid should be all visible
- Check the stability of the set-up, this is crucial for the next step.

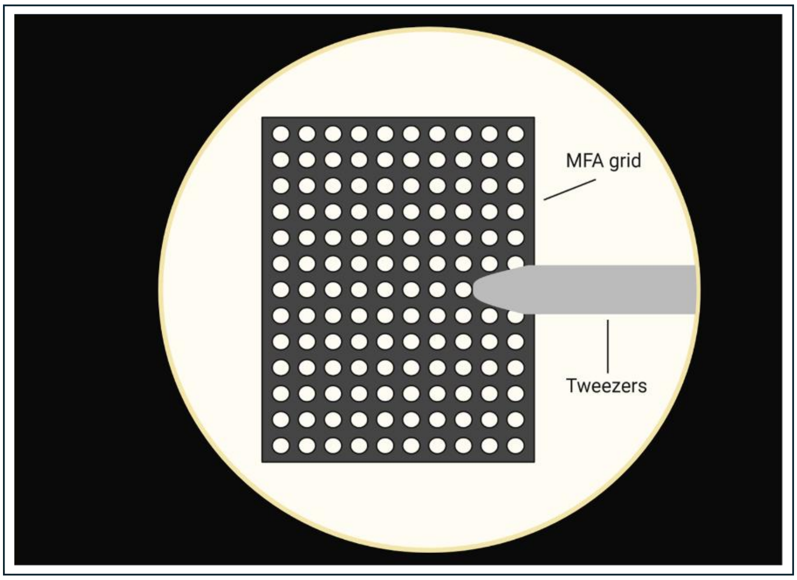

**Figure 2 | MFA Grid Setup.** Schematic representation of the stereoscopic view of the MFA grid prior to fiber insertion. The grid is positioned between tweezers (no. SA13) lined with a thin layer of blu tack. The tweezers are held in place using helping hands. Figure not to scale. Created in bioRender.

### 5. Loading Fibers into the Grid

This section describes how to measure implantation depth, mark fibers accordingly, and insert them into the grid in an organised, damage-minimising sequence. It concludes with securing all implanted fibers in place with adhesive.

#### 5.1 Measuring Implantation Depth

- Using callipers measure out the length of the fiber implantation depth (**Fig. 3**)
    - *Note: refer to the MFA design ensure to factor in the width of the grid (in this case 7 microns).*
- Use a soldering wire to place a drop Norland Optical Adhesive to mark the point of measured depth.
- Treat the adhesive drop with UWAVE curing pen for around 20 seconds (take any necessary precautions when using the UV light).

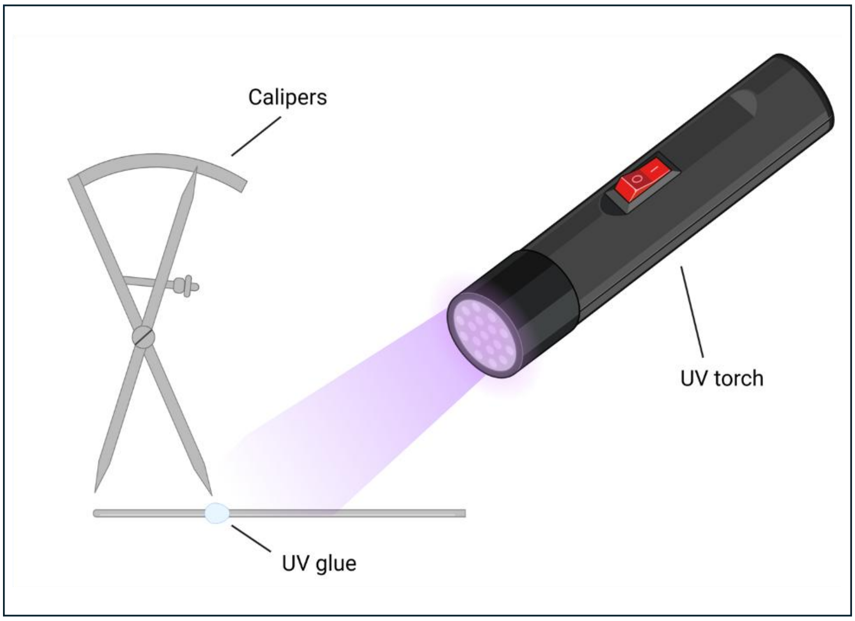

**Figure 3 | Measurement and Marking of Fiber Implantation Depth.** Schematic illustrating the procedure for measuring and marking fiber implantation depths prior to insertion into the MFA grid. Calipers are used to measure the length of the fiber segment to be implanted in reference to the MFA design. The thickness of the grid is included in the final caliper measurement. The measured depth point is marked with UV glue and cured using an UV torch. Figure not to scale. Created in bioRender.

#### 5.2 Insertion Procedure and Securing Fibers in Grid

- In reference to the MFA design, start from the top of the grid and work your way across the rows (this will help avoid the distal ends of the fibers breaking) (**Fig. 4A**).
- To insert the cured fibers, use blue tack lined tweezers no. 5/45 and hold approximately 1/3 from the implantation end to insert cured fibers into assigned grid hole
  - *Note: avoid tapping the end on the grid as this may result in the site being fractured = low efficiency.*
- Once all the fibers have been inserted (**Fig. 4B**), carefully tilt the grid so that it lies perfectly parallel to the workbench.
- Using the soldering wire place a drop of RT80 Black Cyanoacrylate on the top of the grid, the droplet should spread across the grid securing the fibers in place
  - *Optional: use a drop of Rapid Acrylic liquid to speed up the drying process.*

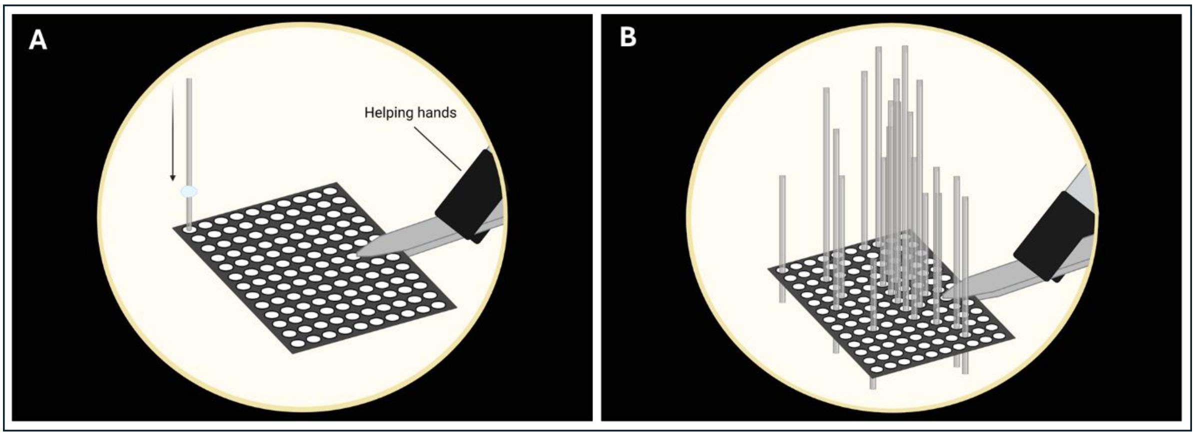

**Figure 4 | Loading Fibers into the MFA Grid.** Schematic representation of the stereomicroscopic view of the fiber loading process. (A) According to the MFA design, cured fibers are inserted into the MFA grid. Beginning at the top of the grid, fibers are inserted row by row (top to bottom) and column by column (left to right). (B) Shows a fully loaded MFA grid**.** Figures not to scale. Created in bioRender.

### 6. Securing and Encasing Distal Fiber Ends

This section describes how to gather, insert, and fix the distal (non-implanting) ends of the fibers inside a protective tube, creating a stable bundle for imaging.

#### 6.1 Tubing Placement

- Cut an 8mm piece using a scalpel Microlumen tubing.
  - *Note: Depending on the number of fibers being implanted choose a suitable size of the Microlumen tubing*
- Bundle the distal (non-implanting) ends of the fibers by gently gathering them with moistened fingertips (**Fig. 5A**)
- Navigate the bundle into the cut Microlumen tubing (**Fig. 5B**).

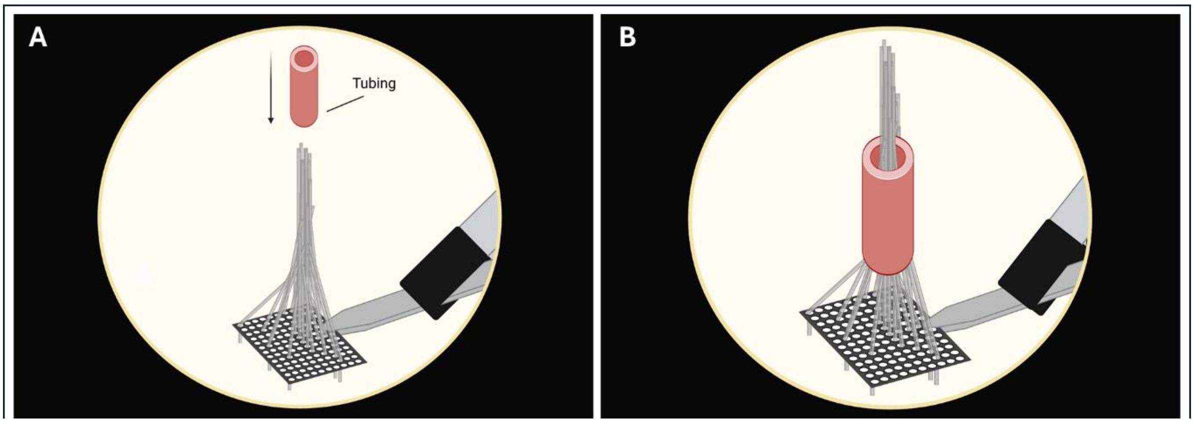

**Figure 5 | Fiber Bundling and Tubing.** Schematic representation of the stereomicroscopic view of fiber bundling and tubing placement process. (A) The distal fiber ends are bundled together using moistened fingertips. (B) Microlumen tubing is then lowered to collect the bundled distal ends. Figures not to scale. Created in bioRender.

#### 6.2 Filling Tube with Adhesive

- Using a micro syringe pull ZAP Thin CA glue, avoiding air bubbles (**Fig. 6**).
- Face the needle towards you, and controlling the decant flow of the glue into the tube fill approximately ¼
- Allow the first layer of glue to dry out (around 10min)
- Once the first layer is dry continue to fill the tube in the same manner, make sure the tube is filled to the brim but not overflowing.
- Leave to dry overnight.
  - *Note: It is key to avoid air bubbles as this will impact the imaging surface*

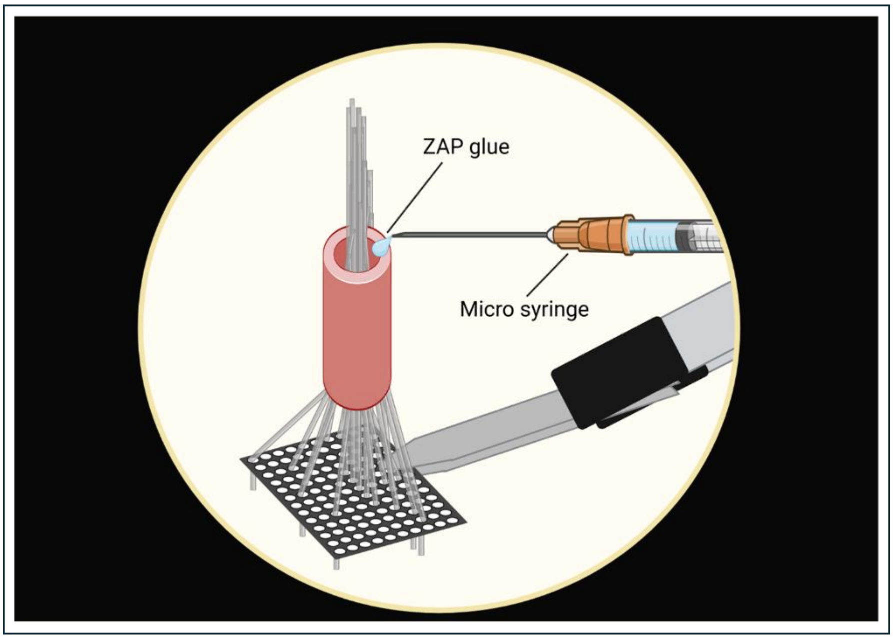

**Figure 6 | Filling Tubing with Adhesive.** Schematic illustrating the process of filling the microlumen tubing with ZAP glue. A micro syringe is used to draw ZAP glue, which is slowly dispensed into the tubing, allowing approximately 10 minutes between layers. This process is repeated until the tube is filled. Figure not to scale. Created in bioRender.

### 7. Encasing MFA and Polishing the Imaging Surface

This section details how to reinforce the MFA with additional adhesive layers and polish the distal surface to ensure a smooth, optically clean imaging area.

#### 7.1 Encasing the grid

- Cut off a short piece of the soldering wire and create a circle
- Slide the Microlumen tube of the MFA halfway into it and tighten the wire, you may wrap the wire around again to ensure stability and twist the ends to secure (**Fig. 7A**).

- Insert the twisted part of the wire into the helping hand and adjust the MFA so that it is parallel to the workbench.
- Use soldering wire to place drops of the RT80 Black Cyanoacrylate to create layers starting from the grid towards the top of the tube. Allow the glue to spread between the fibers (**Fig. 7B**).
- Allow layers to dry out before proceeding with the next one
  - Optional: you can use a drop of Rapid Acrylic liquid to speed the drying process).
- Make sure to cover the red tubing with the black glue to protect the fibers from light and allow to dry

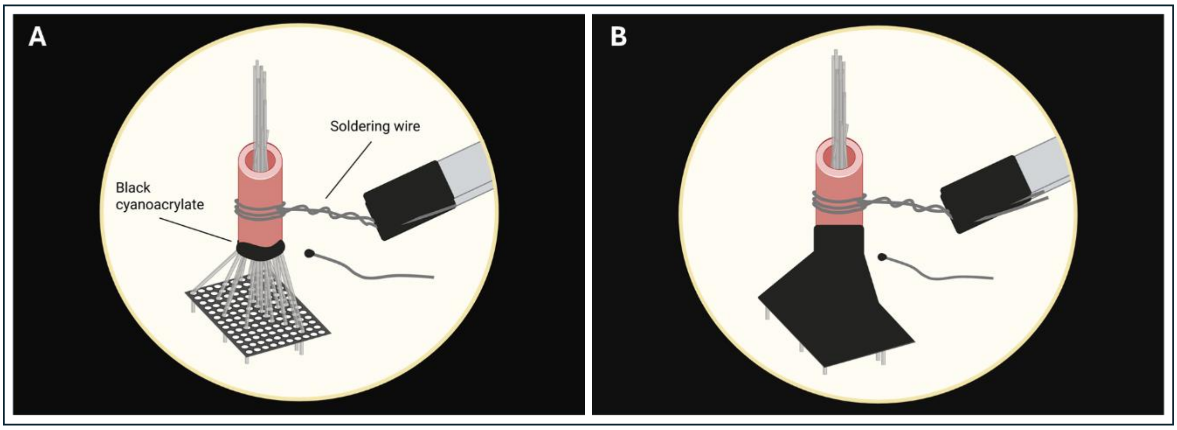

**Figure 7 | Encasing the MFA.** Schematic representation of the stereomicroscopic view of encasing the MFA with black cyanoacrylate. (A) The MFA is repositioned in the helping hands using soldering wire. A segment of soldering wire is dipped in black cyanoacrylate and is used to apply adhesive around the base of the microlumen tubing. (B) A fully encased MFA is shown. Repeat this process, covering the bottom of the microlumen tubing, exposed fibres, and grid is repeated until the MFA is fully encased. Figures not to scale. Created in bioRender.

#### 7.2 Rough Cutting and Polishing Sequence

- Use scissors to trim the distal fiber bundle (you may consider cutting part of the tubing along with it) (**Fig. 8A**).
- Lay out three Thorlabs Diamond Lapping (polishing) sheets of different grit 30μm (green), 6μm (yellow), 3μm (pink).
- Add a drop of distilled water to each of the papers and spread with your finger.
- Grip the tube with tweezers no. 11064-07 and holding the tweezers perpendicular to the paper throughout, using circular motion, polish the surface of the tube for 2 minutes on each of the papers starting with the highest grit (**Fig. 8B**).
- Using the stereoscope examine the imaging surface, ensure the surface is flat and no major air bubbles are obstructing the imaging area.
  - *Note: If the surface still looks uneven you may repolish the tube using the 3μm grit.*

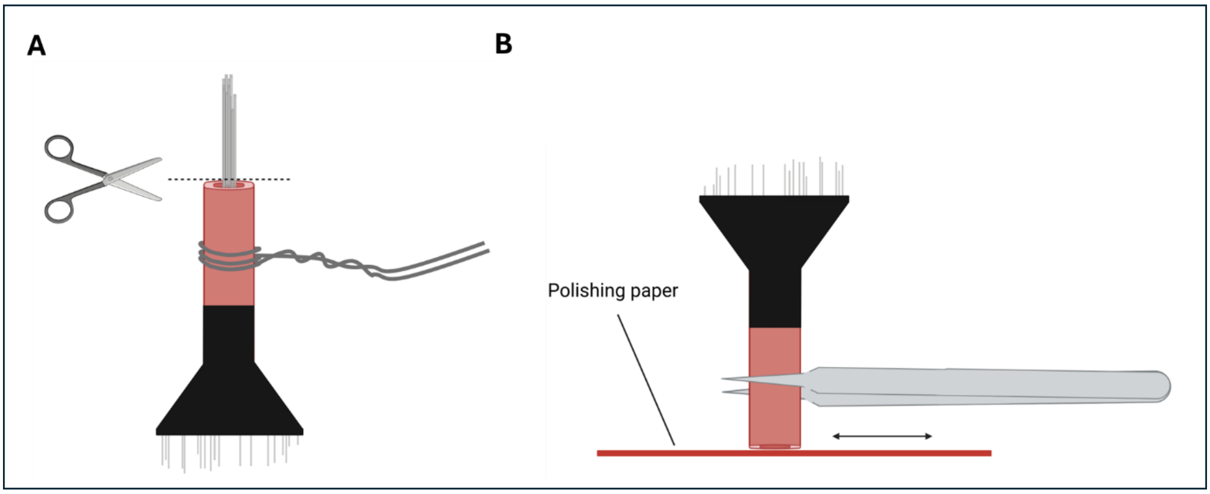

**Figure 8 | Polishing the Imaging Surface.** Schematic of the removal of distal ends and subsequent polishing of the MFA imaging surface. (A) Scissors are used to trim distal fiber ends. (B) The MFA is removed from the soldering wire grip and held with tweezers (no. 11064-07). Holding the MFA perpendicular to the polishing paper, the surface is polished using a back-and-forth or circular motion. This process is repeated for each polishing paper. Figures not to scale. Created in bioRender.

### 8. Fiber Mapping

This section describes how to identify the correspondence between each implanted fiber's grid position and its position on the polished imaging surface. This map is essential for later data interpretation.

#### 8.1 Setup

- Rotate the MFA so that the implantation site is facing upwards and secure in the helping hands by clamping the twisted soldering wire (**Fig. 9**).
- Position the Olympus stereoscope so that you can clearly see the implantation fibers and which row/columns they sit in.
- Turn on the digital Ninyoon WIFI Microscope and place it below the MFA. On your cellular device download the WiFi application (e.g. phone, tablet) you should see the bundled fibers inside the tube (imaging surface).

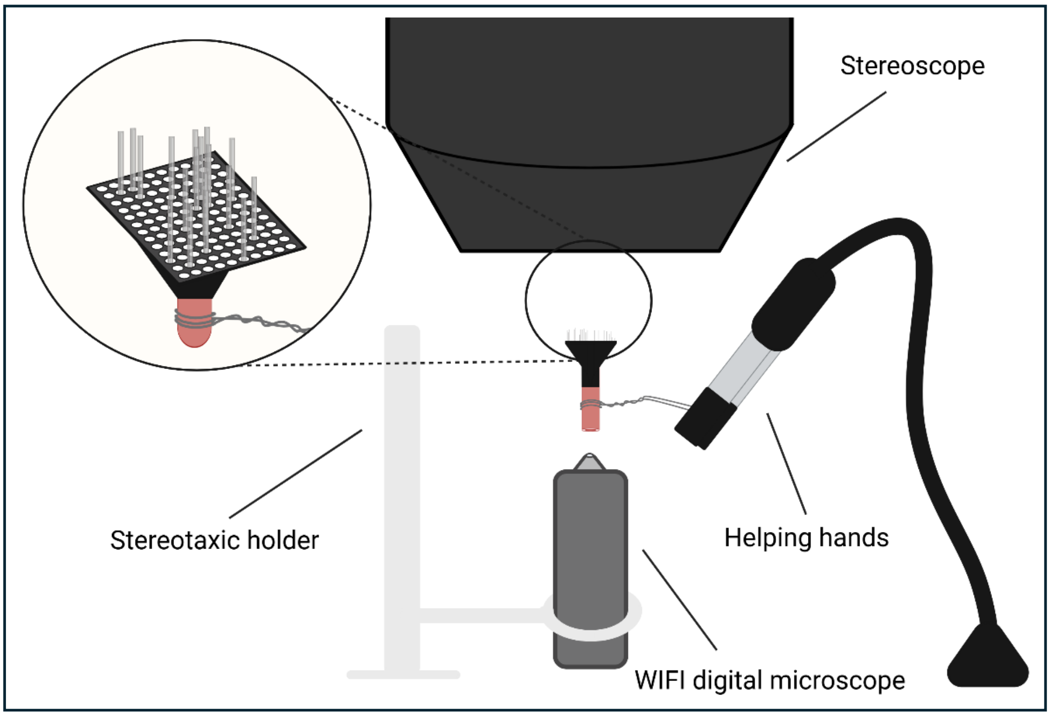

**Figure 9 | MFA Mapping Setup.** Schematic of the preparation and set up for the mapping procedure. The MFA is rotated so that the implantation site is facing upwards towards the stereoscope and is secured using soldering wire and helping hands. A WIFI camera is positioned beneath the imaging surface and is held in place with a stereotaxic holder. The circular inset at the top left shows the stereoscopic view of the setup. Figure not to scale. Created in bioRender.

#### 8.2 Mapping Procedure

- Using a small, 90° folded piece of paper (preferably black) isolate the fibers in the grid row by row going from bottom to top (**Fig. 10A**).
- Shine an LED light pen on the isolated fiber, the fiber should light up on the screen of your cellular device – make a note of which grid coordinate (refer to MFA design) the fiber corresponds to on the imaging surface (keep in mind the image displayed is inverted) (**Figs. 10B&C**).
    - *Note: For rows with multiple fibers, take an additional piece of card to individually isolate the fibers of interest by blocking them from the LED light and following the same process of discrimination.*

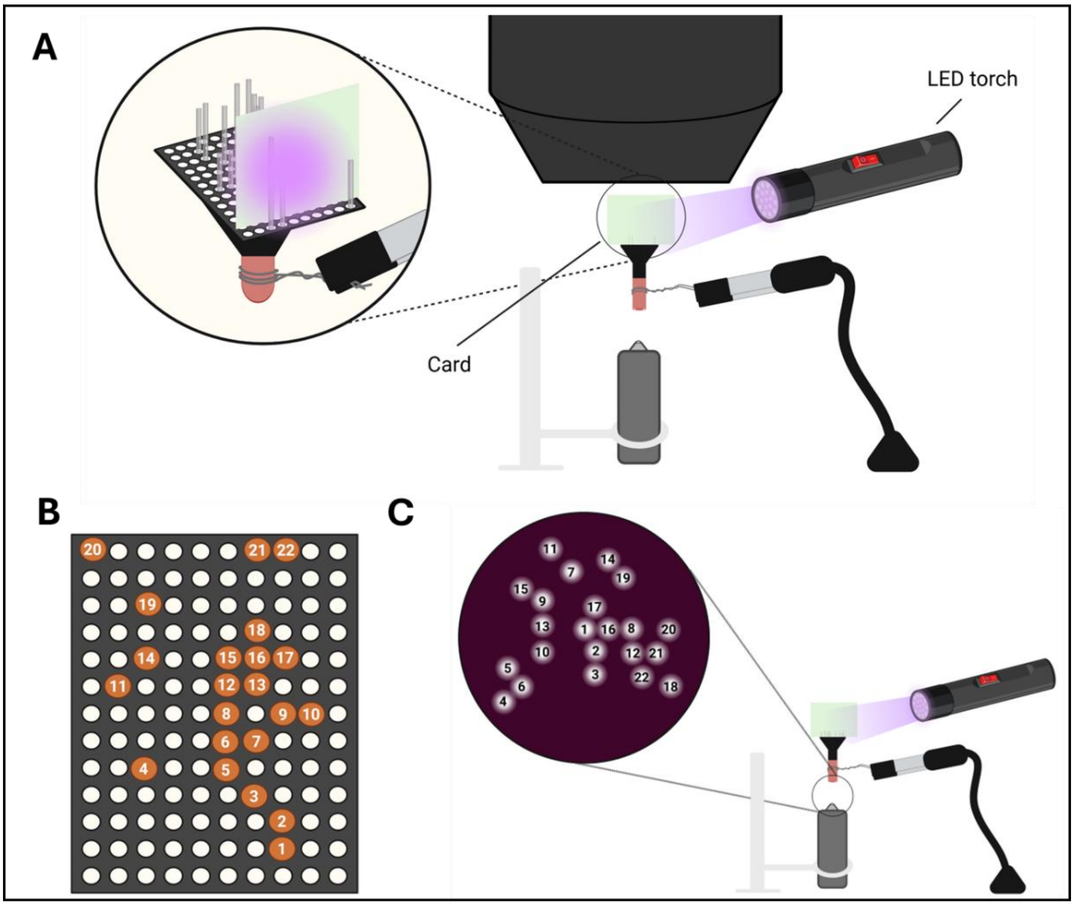

**Figure 10 | MFA Mapping Setup.** Schematic illustrations of the mapping procedure. (A) Setup of the mapping process. Beginning with the bottom row containing a fibers, a piece of card is inserted behind each fiber to isolate it. This is repeated row by row (bottom to top) and column by column (left to right). An LED torch is directed at the isolated fiber, producing illumination visible on the mobile device connected to the WIFI camera. (B) The MFA design, showing the numbering of each fibre according to its layout (bottom to top, left to right. Using this design, the illuminated fiber can be identified on the mobile device screen. (C) Digital microscope setup and inset showing mobile device view of MFA imaging surface with illuminated fibers. The visible fibers are numbered according to the MFA design. Figures not to scale. Created in bioRender.

### 9. Quality Control (QC) and Functional Validation

This section outlines how to validate the functional performance of the MFA using fluorescence testing and image analysis, ensuring the array transmits signals reliably.

#### 9.1 Fluorescence Transmission Testing

- Prepare Fluorescein (TH+) solutions at different concentrations by diluting in PBS; 0µM (PBS), 0.1µM, 0.5µM (optional) and 1µM.
- Place the solutions in the eppendorfs and secure in the eppendorf rack.
- Mount the MFA on the stereotaxic manipulator and lower the implantation ends into the solution starting with the lowest concentration 0µM (PBS).

- Using sCMOS camera connected to the HCImage software observe the imaging surface and capture fluorescence at 405nm (isosbestic) and then at 470nm excitation wavelengths, both on 5V.
    - *Note: Ensure not to change the parameters between capturing, repeat this for each of the concentrations.*

#### 9.2 Data Capture and Analysis

- The captured images should be processed in the MATLAB.
- The analysis should result in a calibration curve, if QC is successful the calibration curve should be linear.
    - *Note: If you have any issues, you may consider repolishing the imaging surface on the lowest 3µm grit again.*
